# Cell-type specific innate immune responses shape rabies virus tropism

**DOI:** 10.1101/2021.07.26.453802

**Authors:** Lena Feige, Tatsuya Kozaki, Guilherme Dias de Melo, Vincent Guillemot, Florence Larrous, Florent Ginhoux, Hervé Bourhy

## Abstract

Viral tropism, or the specificity of a particular virus to infect a certain cell type, is crucial in determining virus replication, viral spread, and ultimately host survival. Rabies, one of the deadliest known zoonotic diseases, is still causing 60.000 human deaths annually. Upon central nervous system (CNS) entry, neurotropic rabies virus (RABV) preserves the neural network by limiting apoptosis and inflammation. To date, we do not fully understand the factors determining RABV tropism and why glial cells are unable to clear RABV from the infected brain. Here, we compare susceptibilities and innate immune responses of CNS cell types towards infection with virulent dog RABV Tha and less virulent Th2P-4M *in vitro*, highlighting differences in cellular susceptibility and antiviral responses. Less virulent Th2P-4M bears mutations introduced in viral phosphoprotein (P-protein) and matrix protein (M-protein) thereby hindering viral immune evasion of the host nuclear factor kappa-light-chain-enhancer of activated B cells (NF-κB) and Janus kinase (JAK) - signal transducer and activator of transcription protein (STAT) pathways. Our results reveal that human neural stem cell (hNSC)-derived neurons and astrocytes, in contrast to human iPSC-derived microglia, are highly susceptible to Tha and Th2P-4M infection *in vitro*. Surprisingly, Th2P-4M presents a stronger neurotropism in hNSC-derived CNS cultures compared to Tha suggesting that NF-κB- and JAK-STAT-mediated antiviral host responses are defining RABV replication and thereby its tropism. Further, we show that astrocyte-like (SVGp12) and microglia-like (HMC3) cells protect neuroblastoma cells (SK-N-SH) from Tha infection *in vitro*. Transcription profiles and quantification of intracellular protein levels revealed major differences in antiviral immune responses mediated by neurons, astrocytes (*IFNB1, CCL5, CXCL10, IL1B, IL6, LIF*), and microglia (*CCL5, CXCL10, ISG15, MX1, IL6*) upon virulent Tha infection. Overall, we provide evidence that RABV tropism depends on its capability to evade cell-type specific immune responses via P- and M-proteins.

**Author summary:** Rabies virus (RABV) neurotropism is widely reported as a unique feature of rabies, and still the exact mechanism underlying RABV susceptibility remains to be elucidated. Several receptors are known to accelerate RABV entry to the cell (1–4) and yet, none of them seems to be essential for successful infection (5,6) questioning a marked cellular tropism. Although RABV is classically reported as strictly neurotropic (7), recent studies report profound infection of glial cells *in vivo* depending on the viral strain and the infection route used (8,9). Here, we provide evidence that human neural stem cell (hNSC)-derived neurons (hiNeurons) and astrocytes (hiAstrocytes) are highly susceptible towards infection with the virulent field RABV strain Tha and less virulent Th2P-4M. In contrast, human iPSC-derived microglia-like cells (hiMicros) are resistant to viral replication *in vitro*. Whereas hiNeurons are immunologically quiescent upon Tha infection, fetal astrocytes and hiMicros establish strong antiviral responses. In contrast to Tha, Th2P-4M, which is unable to evade NF-κB and JAK-STAT pathways (10), shows a more profound neurotropism suggesting that cell-type specific responses shape RABV tropism. Hence, we conclude that viral evasion mechanisms mediated by P- and M-proteins partly determine Tha tropism of human CNS cell types *in vitro*.

## Introduction

Rabies is caused by RABV, a negative-sense single-stranded RNA virus belonging to the family of *Mononegavirales* (11). RABV belongs to the genus *Lyssavirus*, family *Rhabdoviridae*, and presents a negative stranded RNA virus of 12 kb length: the viral genome encodes the nucleoprotein (N-protein), the phosphoprotein (P-protein), the matrix protein (M-protein), the glycoprotein (G-protein), and the RNA-dependent RNA-polymerase (L-protein). RABV, classically referred to as a neurotropic virus, reaches the CNS via retrograde transport along the neural network where it establishes fatal encephalomyelitis in mammals, including humans (12,13). Throughout the whole incubation period, RABV does not cause evident signs of inflammation, complexifying RABV diagnosis (14). Once inside the CNS, RABV successfully hides inside the neural network from glial surveillance (15). Up to date, we do not fully understand the exact mechanisms underlying viral-mediated immune evasion of glial cells.

Viral tropism refers to the ability of a virus to productively infect a certain cell type. Primary infection can be productive, leading to the production of progeny viruses, or abortive, blocking the production of progeny viruses and thereby viral spread. Cellular tropism relies on two major determinants, the expression of entry receptors which enables virus entry and the cellular immune response which allows or restricts productive viral replication (16). The innate immune system therefore represents the first line of defence against viral invaders. It senses viruses via germline-encoded pattern recognition receptors (PRRs) including toll-like receptors (TLRs), retinoic acid-inducible gene I (*RIG-I*) like helicases (RLRs) and nucleotide-binding oligomerization domain-like receptors (NOD-like receptors). The beforehand mentioned receptors recognize viral components, so-called pathogen-associated molecular patterns (PAMPs) or danger-associated molecular patterns (DAMPs). TLR3 and RLRs in the host cells recognize virus-derived RNAs, leading to the activation of the transcription factors, interferon regulatory factor 3 (IRF-3) and nuclear factor κB (NF-κB), which result in the production of type I IFNs and proinflammatory cytokines to establish antiviral responses (17). Type I IFNs then bind type I IFN receptors (IFNAR) and activate the Janus kinase (JAK) and signal transducer and activator of transcription protein 1 (STAT1) signaling pathway, elevating the expression of interferon-stimulated genes (ISGs) with antiviral activity.

RABV uses several receptors to enter cells via clathrin-mediated endocytosis (18–20): nicotinic acetylcholine receptor (nAChR) (1), neuronal cell adhesion molecule (NCAM) (2), low-affinity p75 neurotropin receptor (p75NTR) (3), and metabotropic glutamate receptor subtype 2 (mGluR2) (4). However, those widely expressed receptors are not essential for RABV entry *per se* (21) but lead to an acceleration of RABV infection (6). In contrast to its profound neurotropism *in vivo* (22,23), most cell types are susceptible to RABV infection *in vitro* (24– 26). Whereas most of the research is focussing on the discovery of RABV receptors (1–4) and their interaction with the RABV G-protein (21), less research focusses on how cellular host immune responses shape RABV tropism and how the type of response of the different neural cell types present in the CNS could collectively lead to the establishment of an antiviral response. Recently, several publications reported infection of different glial cells *in vivo*, particularly astrocytes (27,28) and Schwann cells (9), depending on the viral strain and the infection route used (8). Still, we are far from understanding the molecular pathways underlying susceptibility to RABV infection although it is crucial in determining infection outcome.

Here, we investigated cellular immune responses and susceptibilities of different CNS cell types towards Tha and Th2P-4M infection *in vitro*. The RABV field strain Tha was initially isolated from the brain of a Thai patient who died after being bitten by a rabid dog (29). Th2P-4M harbours the same genetic background as Tha, except for mutations introduced into P-(W265G, W287V) and M-proteins (R77K, D100A, A104S, M110L) which were introduced by reverse genetics and which inhibit viral evasion of NF-κB and JAK-STAT pathways (10). We provide evidence that virulent Tha and less virulent Th2P-4M successfully replicate in hiNeurons and hiAstrocytes *in vitro*. In contrast, hiMicros are resistant to virus replication of neither Tha nor Th2P-4M. Surprisingly, Th2P-4M presents a greater neurotropism in hNSC-derived CNS cultures compared to Tha, suggesting that accessory functions of viral P- and M-proteins are determining RABV tropism. Quantification of innate gene expression and protein levels showed that pathogenic Tha is strongly adapted to suppress the innate immune response of neuronal cell types via P- and M-protein-mediated evasion of NF-κB and JAK-STAT pathways, whereas the adaptation mechanisms seem to be less effective for the inhibition of glial immune responses. Whereas successful infection might highly depend on the concentration of RABV receptors on the cellular surface, we suggest that cell type-specific innate immune responses are critical for the success of RABV replication and spread. Hence, our study emphasizes the need to study RABV in relevant CNS culture models to understand the underlying pathways shaping RABV tropism as well as to elucidate the role of glial cells in the RABV-infected brain.

## Results

### Tha and Th2P-4M reveal profound neurotropism in CNS cell lines *in vitro*

Using reverse genetics, we introduced six site-directed mutations in conserved residues of the lyssavirus genus of viral P-protein (30,31) and M-protein (32–34) of the pathogenic dog RABV Tha to diminish RABV pathogenicity (Fig 1A) (10). Upon infection with the two fluorescent RABV strains, Tha-eGFP and Th2P-4M-eGFP respectively, eGFP was expressed in all different CNS cell lines indicating successful infection of neuronal SK-N-SH, astrocyte-like SVGp12, and microglia-like HMC3 cells *in vitro* (Fig 1B). Quantification of eGFP expression via fluorescence microscopy revealed that, in average, 78.87% of SK-N-SH (95% confidence interval (CI_95_) 73.1 – 84.65%) expressed eGFP whereas only 21.26% of astrocyte-like SVGp12 (CI_95_ 18.95 – 23.58%) and 25.43% of microglial-like HMC3 (CI_95_ 23.15 – 27.71%) cells expressed eGFP upon Tha-eGFP infection (Fig 1C). In contrast to SVGp12 and HMC3, significantly less SK-N-SH (CI_95_ 60.19 – 68.5%) expressed eGFP upon Th2P-4M infection compared to Tha-eGFP (p-value<0.0001). Both viral strains revealed a profound neurotropism since Th2P-4M-eGFP, like Tha-eGFP, induced eGFP expression in significantly less astrocyte-like SVGp12 (CI_95_ 15.74 – 19.46%) and microglial-like HMC3 cells (CI_95_ 20.5 – 23.12%) compared to Tha-eGFP-infected SK-N-SH (p-value<0.0001).

**Fig 1.**
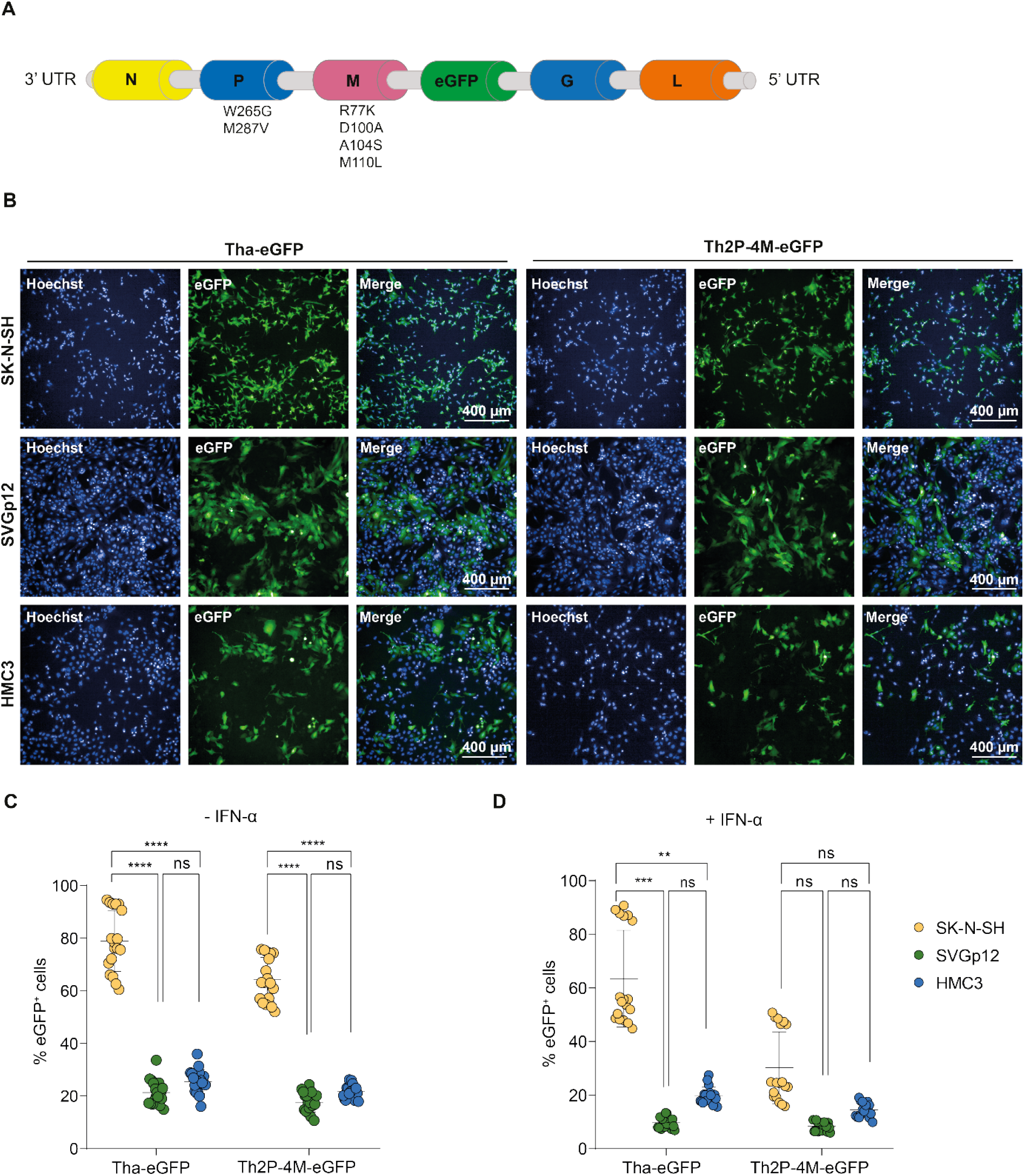
Tha-eGFP and Th2P-4M-eGFP preferentially replicate and spread in human neuroblastoma cells (SK-N-SH) compared to astrocyte-like (SVGp12) and microglia-like (HMC3) cell lines *in vitro*. (A) Schematic overview of the genomic organization of Tha-eGFP and Th2P-4M-eGFP. Tha-eGFP and Th2P-4M-eGFP harbour the same genetic background except for two mutations introduced in viral P-protein (W265G, W287V) and four mutations introduced in the M protein (R77K, D100A, A104S, M110L) (10,31,32). For imaging purposes, the eGFP sequence was introduced after the M protein gene sequence as described previously (35). (B) Representative immunofluorescence pictures of SK-N-SH, SVGp12, and HMC3 cells upon infection with Tha-eGFP and Th2P-4M-eGFP. (C) Quantification of eGFP^+^ cells in human CNS monocultures upon Tha-eGFP or Th2P-4M-eGFP infection. (D) Quantification of eGFP^+^ cells in human CNS monocultures upon Tha-eGFP or Th2P-4M-eGFP infection and subsequent IFN-α treatment at 24 hours post-infection. (B-D) Cells were infected with Tha-eGFP or Th2P-4M-eGFP (MOI 0.5) and imaged at 48 hours post-infection. All experiments were performed three times (n=3) independently. (C-D) Each dot represents imaging of one well of a 96-well-plate (approx. 8×10^3^ cells/well). Bars show mean ± SD with a Tukey’s multiple comparisons test (**** adjusted p-value<0.0001, *** adjusted p-value<0.001, ** adjusted p-value<0.01, * adjusted p-value<0.05). eGFP = Enhanced Green Fluorescent Protein.

Adding exogenous IFN-α artificially activates the JAK-STAT pathway resulting in the expression of ISGs and antiviral proteins. Here, exogenous IFN-α significantly lowered virulent Tha-eGFP infection of SK-N-SH to 63.45% (CI_95_ 54.48 – 72.42%, p-value<0.0001) and Th2P-4M-eGFP infection was reduced to 30.16% (CI_95_ 23.45 – 36.82%, p-value<0.0001) compared to infected non-treated SK-N-SH. Similarly, exogenous IFN-α lowered resulted in significantly less SVGp12 cells expressing eGFP upon pathogenic Tha-eGFP infection (CI_95_ 8.66 – 10.67%, p-value=0.0004) and less-pathogenic Th2P-4M-eGFP infection (CI_95_ 7.55 – 9.21%, p-value<0.0105). In contrast, adding IFN-α to HMC3 did not result in a significant reduction in eGFP^+^ cells upon Tha-eGFP (CI_95_ 18.27 – 21.36%, p-value=0.3083) or Th2P-4M-eGFP infection (CI_95_ 13.17 – 15.8%, p-value=0.9801, Fig 1D). Comparing IFN-α-mediated effects in inhibiting viral replication revealed the most striking differences between Tha-eGFP and Th2P-4M-eGFP-infected SK-N-SH: exogenous IFN-α lowered Tha-eGFP infection only by 19.55% whereas Th2P-4M-eGFP infection was lowered by 53.13%. Thus, conserved residues in viral P-protein and M-protein of Tha-eGFP seem to be crucial to evade the IFN-α-induced antiviral responses of neuronal cells.

### Glial cell lines constitutively protect SK-N-SH from RABV infection

Since CNS monocultures do not reflect the complexity of the CNS, we investigated if glial cells limit RABV infection in RABV-infected CNS co-culture models. Therefore, we compared the percentage of infection of neuroblastoma monocultures (SK-N-SH) to co-cultures of neuroblastoma and astrocyte-like cells (SK-N-SH + SVGp12), co-cultures of neuroblastoma and microglia-like cells (SK-N-SH + HMC3) and co-cultures of neuroblastoma, astrocyte- and microglia-like cells (SK-N-SH + SVGp12 + HMC3, Fig 2A). During Tha-eGFP and Th2P-4M-eGFP infection, the presence of SVGp12, HMC3, or both cell types in co-cultures significantly reduced the percentage of eGFP^+^ cells compared to SK-N-SH monocultures (Fig 2A). Nevertheless, the presence of glial cells lowered the percentage of eGFP^+^ cells in Tha-eGFP-infected co-cultures not more than expected beforehand from lower susceptibilities observed in Tha-eGFP-infected glial monocultures (Fig 1C, S1 Fig). In contrast, the presence of glial cells (SVGp12, HMC3, or SVGp12 and HMC3) significantly reduced the percentage of eGFP^+^ cells more than expected beforehand from lower susceptibilities observed in Th2P-4M-eGFP-infected glial monocultures (Fig 2A, S1 Fig). Therefore, our data suggest that Th2P-4M-eGFP induces the secretion of antiviral proteins which lead to a crosstalk between SK-N-SH, astrocyte-like SVGp12, and microglia-like HMC3 cells which in turn enhances restriction of viral replication compared to the wildtype infection.

**Fig 2.**
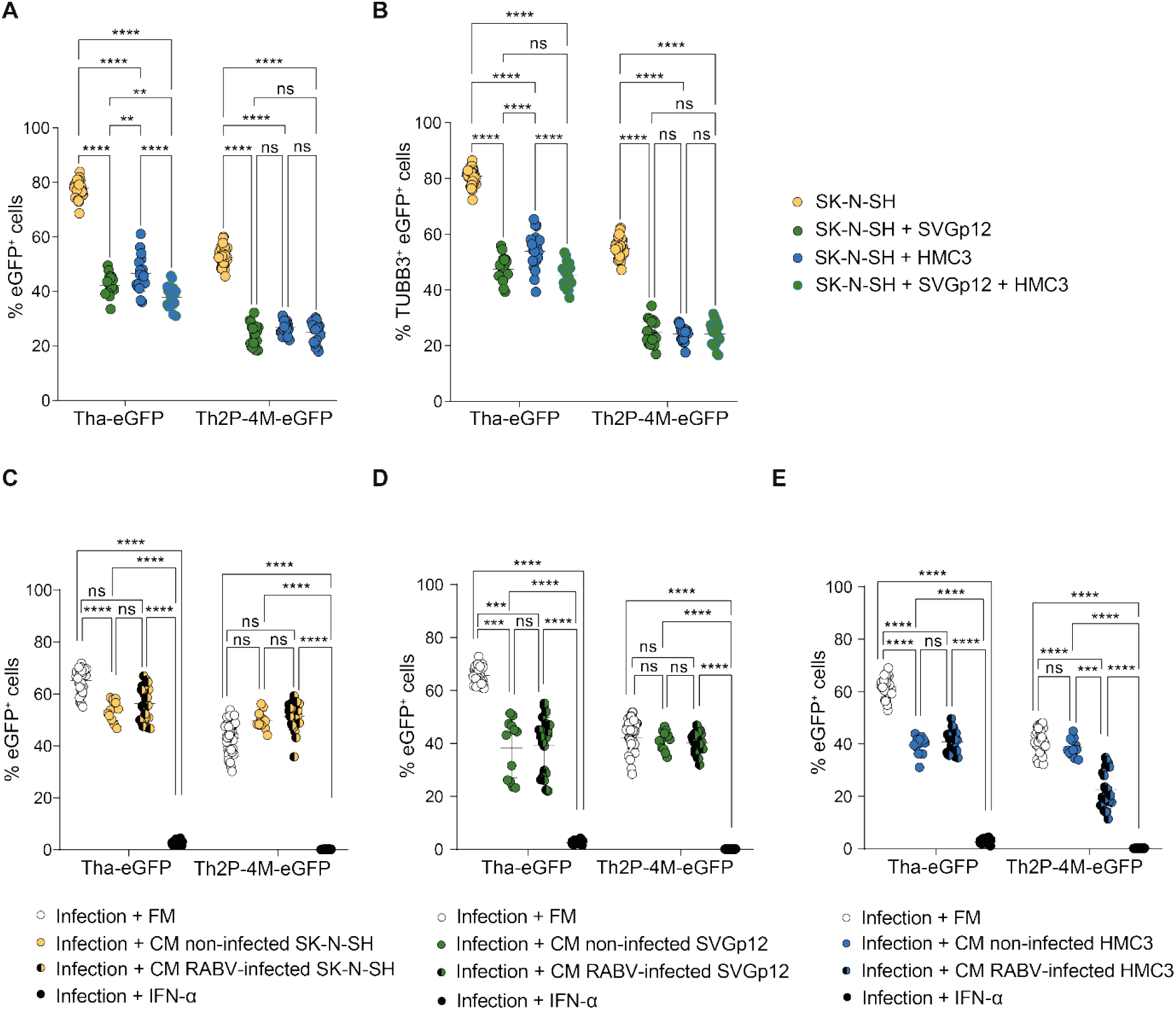
**Astrocyte-like SVGp12 and microglia-like HMC3 protect neuroblastoma cells from Tha-eGFP infection *in vitro*. (A) Quantification of eGFP^+^ cells in mono- and co-cultures. (B) Quantification of TUBB3**^**+**^ **eGFP**^**+**^ **cells in mono- and co-cultures**. (A – B) Mono- or co-cultures were infected with Tha-eGFP or Th2P-4M-eGFP (MOI 0.5) and imaged at 48 hours post-infection using the OPERA Phenix^®^ High Content Screening System (Perkin Elmer). (C-E) Quantification of eGFP^+^ cells in SK-N-SH monocultures. After infection with Tha-eGFP or Th2P-4M-eGFP (MOI 0.5), SK-N-SH cells were incubated with the filtered fresh medium (FM) or filtered conditioned medium (CM) from non-infected or homotypic infected (MOI 5, 24 hours post-infection) SK-N-SH (C), SVGp12 (D), or HMC3 cells (E). In detail, THA-eGFP-infected cells were only treated with supernatants from cells previously infected with THA-eGFP. Correspondingly, TH2P-4M-eGFP-infected cells were only treated with supernatants from cells previously infected with TH2P-4M-eGFP. Cells were imaged at 48 hours post-infection. (A – E) All experiments were performed three times (n=3) independently. Each dot represents imaging of one well of a 96-well-plate (approx. 8×10^3^ cells/well). Bars show mean ± SD with a Tukey’s multiple comparisons test (**** adjusted p-value<0.0001, *** adjusted p-value<0.001, ** adjusted p-value<0.01, * adjusted p-value<0.05). CM = conditioned medium; eGFP = Enhanced Green Fluorescent Protein; FM = filtered fresh medium; TUBB3 = Class III Beta-Tubulin.

Since RABV is a neurotropic virus that enters the CNS via the neural network, we explored if the presence of glial cells is restricting RABV infection of SK-N-SH in CNS co-culture models (Fig 2B). Cells of glial origin, SVGp12 or HMC3, significantly reduced infection of Class III Beta-Tubulin (TUBB3^+^) neuronal cells during Tha-eGFP and Th2P-4M-eGFP infection (Fig 2B). During Tha-eGFP infection, the presence of astrocyte-like SVGp12 lowered the percentage of TUBB3^+^ eGFP^+^ cells stronger compared to the presence of microglia-like HMC3. Adding both, SVGp12 and HMC3, lowered the percentage of TUBB3^+^ eGFP^+^ cells in Tha-eGFP-infected co-cultures the most suggesting a synergistic effect. Unlike Tha-eGFP, the presence of SVGp12, HMC3, or both cell types together lowered the percentage of TUBB3^+^ eGFP^+^ cells in Th2P-4M-eGFP-infected co-cultures equally (Fig 2B).

Further, we investigated if glial-mediated restriction of SK-N-SH infection is mediated via the secretion of glial-derived proteins to the supernatant. Therefore, we transferred the medium of non-infected or homotypic infected SK-N-SH, SVGp12 or HMC3 cells after viral particle-removal, to freshly infected SK-N-SH cells to see if conditioned medium reduces the percentage of eGFP^+^ SK-N-SH compared to the transfer of fresh cell culture medium (Fig 2C-E). During Tha-eGFP infection, transfer of conditioned medium originating from non-infected SK-N-SH to Tha-eGFP-infected SK-N-SH cells lowered the percentage of eGFP^+^ SK-N-SH significantly (p-value<0.0001). Surprisingly, transfer of conditioned medium originating from Tha-eGFP-infected SK-N-SH cells did not result in a significant reduction of eGFP^+^ SK-N-SH (p-value=0.40; Fig 2C), suggesting that Tha-eGFP is blocking the basal secretion of antiviral mediators in infected SK-N-SH cells compared to non-infected SK-N-SH. In contrast to SK-N-SH (Fig 2C), transfer of medium from non-infected as well as Tha-eGFP-infected SVGp12 (p-value<0.0001) and HMC3 (p-value<0.0001) significantly reduced the percentage of eGFP^+^ SK-N-SH (Fig 2D-E).

During Th2P-4M-eGFP infection, transfer of non-infected supernatant from SK-N-SH, SVGp12, or HMC3 did not reduce the percentage of eGFP^+^ SK-N-SH (Fig 2C-E). Whereas neither transfer of Th2P-4M-eGFP-infected SK-N-SH (Fig 2C) nor Th2P-4M-eGFP-infected SVGp12 (Fig 2D) lowered the percentage of eGFP^+^ SK-N-SH, Th2P-4M-eGFP-infected HMC3 significantly reduced the percentage of eGFP^+^ SK-N-SH (p-value<0.0001; Fig 2E). This leads to the hypothesis that Th2P-4M induces the secretion of antiviral proteins in HMC3 which in turn restrict Th2P-4M-eGFP replication in SK-N-SH. During both Tha-eGFP and Th2P-4M-eGFP infection of SK-N-SH, adding exogenous IFN-α strongly restricted viral infection (p-value<0.0001; Fig 2C-E).

Overall, non-infected SK-N-SH, SVGp12 and HMC3 constitutively protect SK-N-SH from virulent Tha-eGFP infection via the secretion of proteins. In contrast to Tha-eGFP-infected SK-N-SH, however, glial cells protect SK-N-SH also upon virulent Tha-eGFP infection (Fig 2C-E).

### RABV infects human NSC-derived hiNeurons and hiAstrocytes but fails to infect hiMicros *in vitro*

After investigating the role of astrocyte-like and microglia-like cells in restricting RABV infection, we investigated the role of human stem cell-derived glial cells on RABV infection. First, we differentiated hNSC into hiNeurons and hiAstrocytes and iPSC into hiMicros according to a published protocol (36). To obtain hiMicros, we differentiated iPSC-derived hiMacs for three consecutive weeks with hiNeurons and hiAstrocytes co-cultures. Prior to infection, cultures revealed in average 71.0% hiNeurons (CI_95_ 69.92-72.05%), 18.4% hiAstrocytes (CI_95_ 17.24-19.61%), and 10.6% hiMicros (CI_95_ 9.03-12.15%, S2 Table). After differentiation, cellular fate was validated by immunofluorescence (Fig 3A), RT-qPCR, or FACS (S2 Fig). Upon infection, Tha-eGFP and Th2P-4M-eGFP successfully replicated in hiNeurons and hiAstrocytes but failed to infect hiMicros (Fig 3A). Since differentiation of hNSC to hiNeurons resulted in a culture containing in average 85.6 % hiNeurons (CI_95_ 84.31-86.02%, S1 Table) but also not negligible proportions of hiAstrocytes (CI_95_ 13.98-15.69%, S1 Table), we investigated viral tropism in co-cultures containing hiNeurons and hiAstrocytes and co-cultures containing hiNeurons, hiAstrocytes, and hiMicros. Upon infection, both viral strains showed a neurotropic behaviour in hiNeurons and hiAstrocytes co-cultures (Fig 3B). In contrast to CNS cell line co-cultures (Fig 2A), Th2P-4M-eGFP revealed a more pronounced neurotropism in hNSC-derived co-cultures compared to Tha-eGFP (Fig 3B).

**Fig 3.**
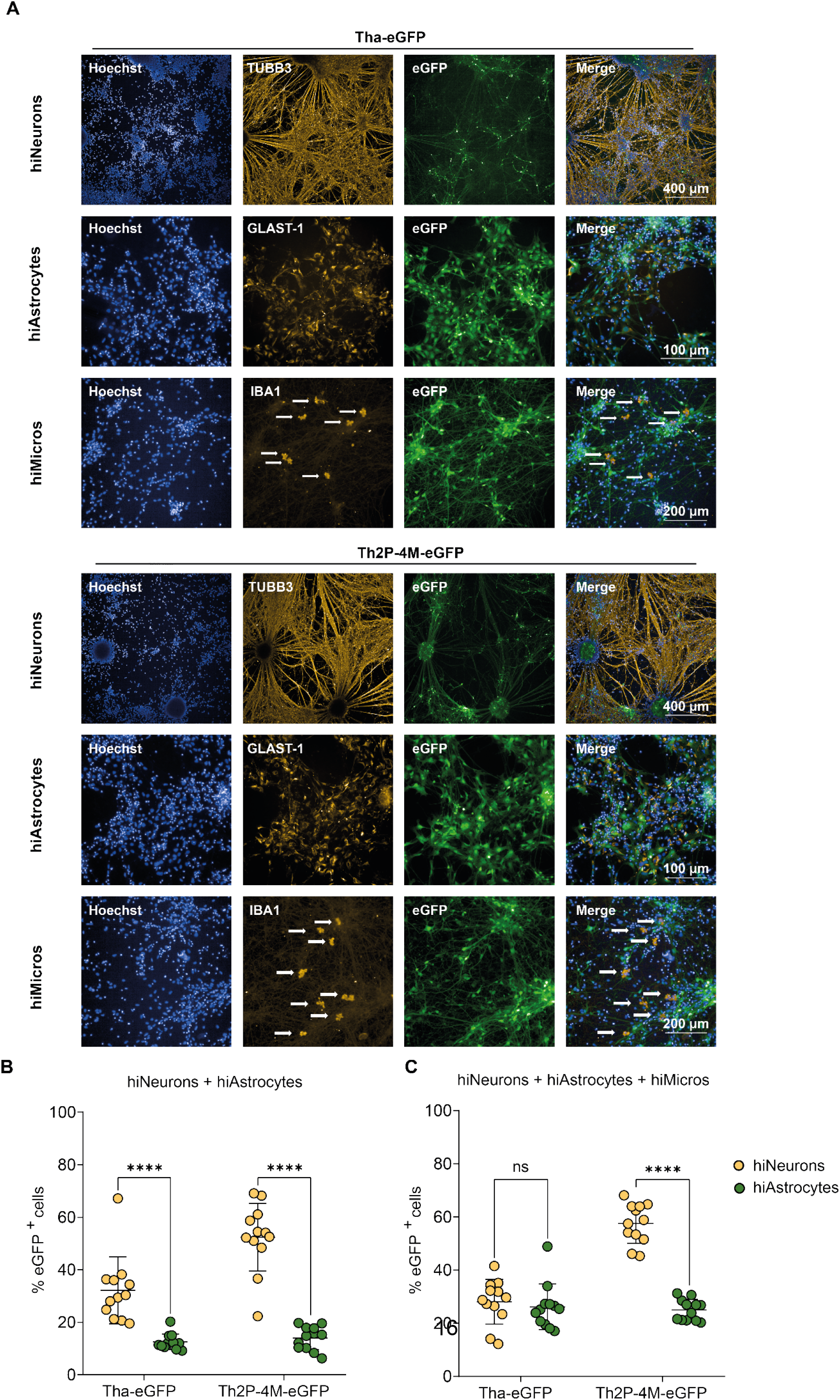
Whereas hiNeurons and hiAstrocytes are highly susceptible towards Tha-eGFP and Th2P-4M-eGFP infection, hiMicros are resistant to Tha-eGFP or Th2P-4M-eGFP infection *in vitro*. (A) Representative immunofluorescence pictures of hiNeurons, hiAstrocytes and hiMicros upon infection with Tha-eGFP and Th2P-4M-eGFP. Cells were stained in cultures consisting of hiNeurons, hiAstrocytes, and hiMicros (S2 Table). (B) Quantification of eGFP^+^ cells in co-cultures consisting of hiNeurons and hiAstrocytes. Prior to infection, cultures consisted of 85.6% hiNeurons and 14.4% hiAstrocytes (S1 Table). (C) Quantification of eGFP^+^ cells in co-cultures consisting of hiNeurons, hiAstrocytes and hiMicros. Prior to infection, cultures consisted of 71.0% hiNeurons, 18.4% hiAstrocytes, and 10.6% hiMicros (S2 Table). (A-C) Cells were infected with Tha-eGFP or Th2P-4M-eGFP (MOI 0.5) and imaged at 48 hours post-infection. All experiments were performed three times (n=3) independently. (B-C) Every dot represents imaging of one well of a 96-well-plate (approx. 1×10^4^ cells/well). Bars show mean ± SD with an unpaired two-tailed t-test (****p-value<0.0001, *** p-value<0.001, ** p-value<0.01, * p-value<0.05). eGFP = Enhanced Green Fluorescent Protein; GLAST-1 = Glutamate Aspartate Transporter-1; IBA1 = Ionized calcium-binding adaptor molecule 1; TUBB3 = Class III Beta-Tubulin.

Further, we wanted to compare the influence of hiMicros on RABV tropism. Adding hiMicros to hiNeurons and hiAstrocytes co-cultures influenced RABV tropism and limited Tha-eGFP neurotropism *in vitro*: whereas Tha-eGFP infected 28.07% hiNeurons (CI_95_ 22.73 – 33.42%) and 26.15% hiAstrocytes (CI_95_ 20.71 – 31.58%), Th2P-4M-eGFP infected 57.49% hiNeurons and 25% hiAstrocytes (CI_95_ 22.46 – 27.54%, Fig 3C). In contrast to Th2P-4M-eGFP, adding hiMicros to the co-cultures reduced the neurotropism of Tha-eGFP. Despite controversial results obtained from cell line co-cultures and stem cell-derived co-cultures, both models indicate an influence of glial cells in shaping RABV neurotropism.

### Virulent Tha strongly suppresses the innate immune gene expression of neural cell types *in vitro*

To further investigate the roles of glial cells in inducing cell-specific immune responses, different CNS cell types were cultured in monocultures and the expression of a panel of selected innate immunity genes (*TLR3, TLR7, IFIH1, DDX58, DHX58, IRF7, IFNB1, CCL5, CXCL10, ISG15, MX1, IL1B, IL6, LIF*) was quantified via RT-qPCR.

Comparing basal expression levels of innate immunity genes between CNS cell types by RT-qPCR revealed that hiMicros present a strong basal expression of RLRs (*IFIH1, DDX58, DHX58*) *TLR3*, adaptor molecules (*IRF7*), chemokines (*CCL5*) and antiviral proteins (*MX1*). In contrast, fetal pAstrocytes revealed a strong expression of the RLR *DDX58* (encoding RIG-I) and SK-N-SH showed a strong expression of *TLR3*, although this was not observed in hiNeurons (S3A Fig). Upon IFN-α treatment, astrocytes, astrocyte-like cells, microglia-like, and neuroblastoma cells strongly amplified the inflammatory response via the expression of *TLR3, IFIH1, DDX58, DHX58, IRF7, ISG15, and MX1*. In contrast, hiNeurons showed little response to IFN-α stimulation. In detail, *DDX58, ISG15* and *MX1* were the only genes that were modestly modulated by IFN-α in hiNeurons (S3B Fig). Additionally, we also noticed a strong and specific amplification of *CXCL10* in fetal pAstrocytes upon IFN-α treatment.

In the next step, we quantified the expression of innate immunity genes upon Tha and Th2P-4M infection via RT-qPCR (Fig 4A and 4B): first, Th2P-4M induced a stronger innate immune response compared to Tha. Despite the induction of strong immune responses, neither Tha nor Th2P-4M strongly induced cell death or apoptosis in neuroblastoma, astrocyte-like or microglia-like cells (S4 Fig). Secondly, fetal pAstrocytes and hiMicros strongly induced the expression of innate immunity genes upon Tha and Th2P-4M infection which was not observed in respective astrocyte-like (SVGp12) and microglia-like (HMC3) cell lines. Lastly, little differences were observed between CNS cell lines (SK-N-SH, SVGp12, and HMC3), whereas profound differences were observed in the innate immune response of hiNeurons, fetal pAstrocytes, and hiMicros (S5 and S6 Fig). Upon Tha and Th2P-4M infection, fetal pAstrocytes strongly induced the expression of *IFIH1, CCL5*, and *CXCL10* whereas hiMicros strongly upregulated expression of *CXCL10* and genes coding for antiviral proteins *ISG15* and *MX1* (Fig 4A).

**Fig 4.**
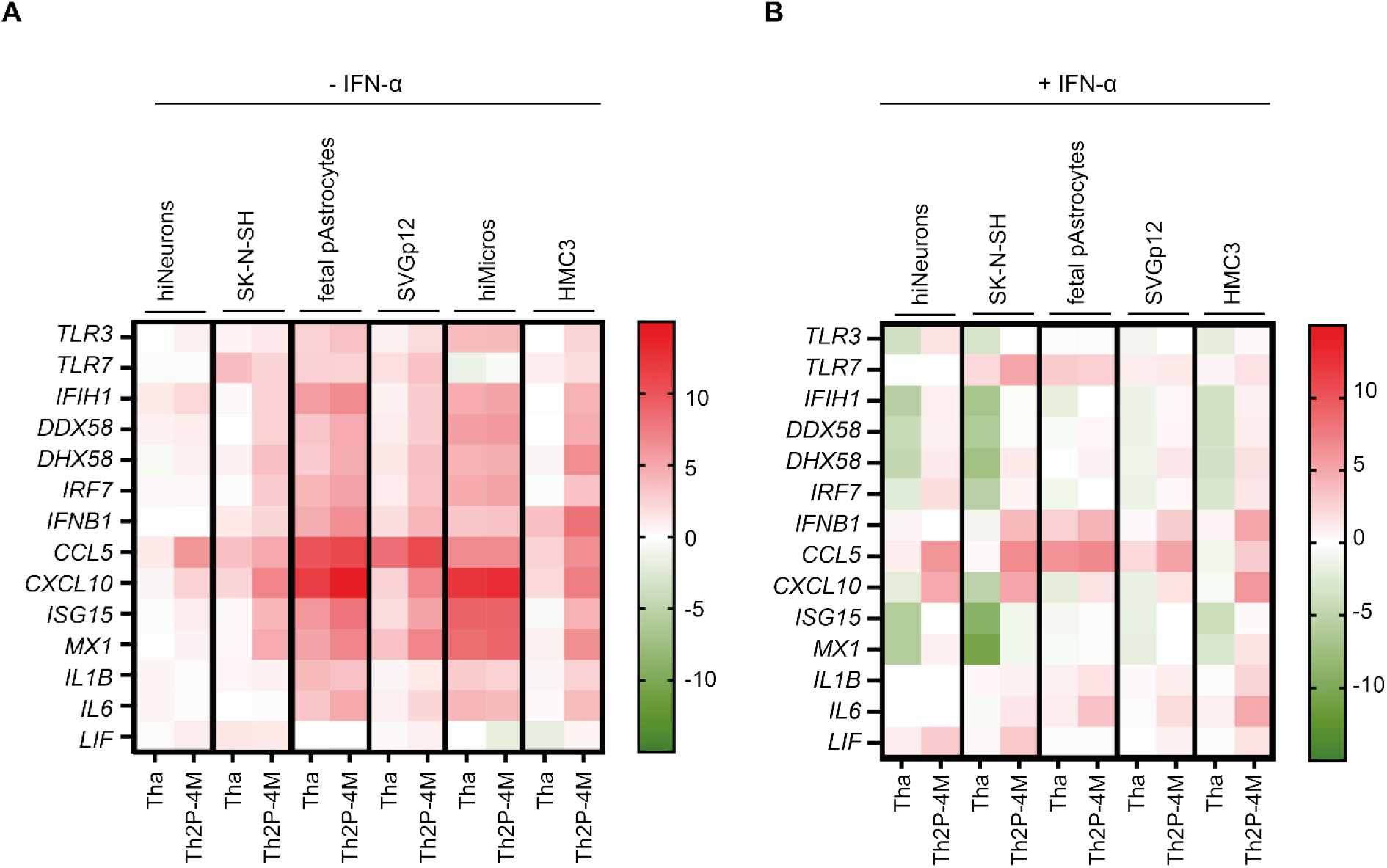
The virulent RABV strain Tha is highly adapted to evade the neural innate immune system. (A) Expression of innate immunity genes upon Tha and Th2P-4M infection. (B) Expression of innate immunity genes upon Tha and Th2P-4M infection with subsequent IFN-α treatment at 24 hours post-infection. (A-B) Cells were infected with Tha or Th2P-4M (MOI 5) and gene expression was quantified at 48 hours post-infection via RT-qPCR. To obtain hiNeurons, hNSC were differentiated for 21 days resulting in cultures which contained 85.6% hiNeurons (S1 Table). hiMacs were differentiated to hiMicros by co-culturing them for three consecutive weeks in inserts with hiNeurons (see Material and Methods). All experiments were performed three times (n=3) independently. Heatmaps present gene expression (ΔΔCT) which was normalized to the expression of the reference gene *18S* and the respective mock-infected (A) or IFN-α-treated mock (B). Colour scaling presents differences observed in gene expression (ΔΔCT) between infected and respective non-infected controls.

Subsequently, we stimulated cells with IFN-α at 24 hours post-infection to artificially activate the JAK-STAT pathway. Expression of innate immunity genes was quantified at 48 hours post-infection to test if Tha or Th2P-4M can inhibit the IFN-α-induced inflammatory response (Fig 4B). Profound differences were observed without IFN-α treatment (Fig 4A), whereas cell type-specific differences were less profound after IFN-α stimulation (Fig 4B, S5 and S6 Fig). Despite IFN-α treatment, virulent Tha suppressed the expression of innate immunity genes compared to less virulent Th2P-4M. In detail, IFN-α-treated fetal pAstrocytes strongly expressed *TLR7, IFNB1, CCL5, IL1B*, and *IL6* upon Tha infection. IFN-α-treated microglia-like cells upregulated *IFNB1, CCL5, CXCL10*, and *IL6* upon Tha infection (Fig 4B). In contrast to Th2P-4M, Tha strongly downregulated the expression of *TLR3*, RLRs (*IFIH1, DDX58, DHX58*) and the antiviral proteins *MX1* and *ISG15* in IFN-α-treated cells of neuronal origin (SK-N-SH and hiNeurons) as well as expression of RLRs (*IFIH1, DDX58, DHX58*) and *ISG15* in microglia-like HMC3.

Overall, we show that virulent Tha induces little innate immune responses in cells of neuronal origin (SK-N-SH and hiNeurons), whereas glial cells (fetal pAstrocytes and hiMicros) strongly respond to virulent Tha infection (Fig 4, S5 and S6 Fig).

### Glial cells produce IL-1β and IL-6 upon Tha and Th2P-4M infection

To see if cell-specific immune responses can be observed by quantifying protein expression levels of infected cells, we quantified human IFN-β, IFN-γ, IL-1β, IL-6, IL-15, CXCL10, LIF, CCL5, and TNF-α protein expression intracellularly (Fig 5). Intracellular protein concentrations revealed the induction of cell-specific immune responses upon RABV infection: both neuronal cell types investigated, SK-N-SH and hiNeurons, did not induce any of the beforementioned proteins upon Tha or Th2P-4M infection but constitutively expressed modest levels of TNF-α (Fig 5A-B). In contrast, Tha and Th2P-4M infection induced expression of IL-1β and LIF in astrocyte-like SVGp12 although observed differences were not significant (Fig 5C). Pure cultures of fetal pAstrocytes did not induce the expression of any of the abovementioned proteins upon neither Tha nor Th2P-4M infection but constitutively expressed modest levels of LIF and TNF-α (Fig 5D). Microglia-like HMC3 induced expression of IL-6 upon Tha and Th2P-4M infection. Further HMC3 constitutively expressed moderate levels of LIF and TNF-α (Fig 5E).

**Fig 5.**
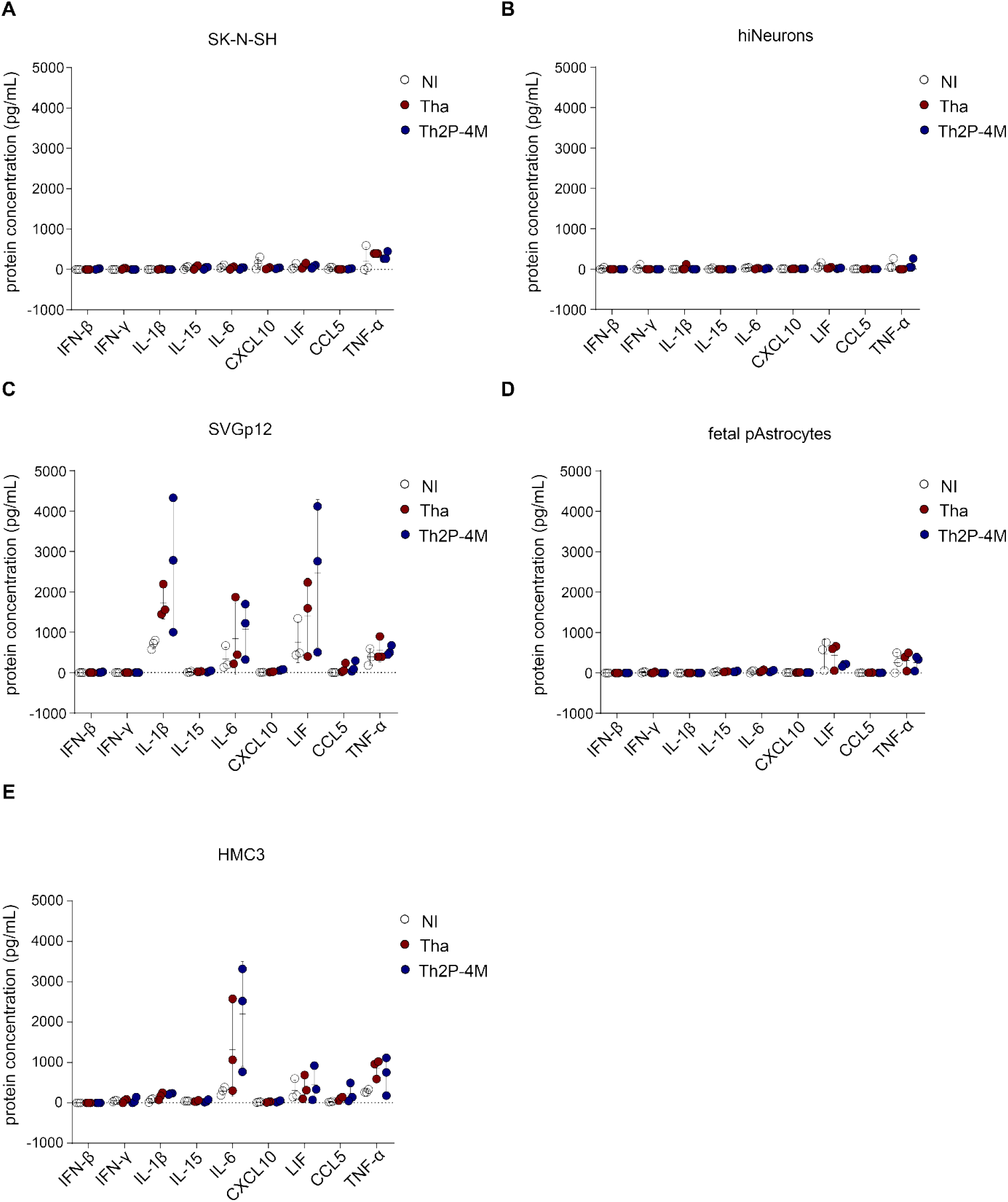
Astrocytes, astrocyte-like SVGp12 and microglia-like cells exhibit a cell-specific induction of protein expression upon Tha and Th2P-4M infection. (A) Intracellular protein concentrations of SK-N-SH cells upon Tha or Th2P-4M infection. (B) Intracellular protein concentrations of hiNeurons upon Tha or Th2P-4M infection. (C) Intracellular protein concentrations of astrocyte-like SVGp12 cells upon Tha or Th2P-4M infection. (D) Intracellular protein concentrations of fetal pAstrocytes upon Tha or Th2P-4M infection. (E) Intracellular protein concentrations of microglia-like HMC3 cells upon Tha or Th2P-4M infection. (A-E) Cells were infected with Tha or Th2P-4M (MOI 5) and intracellular protein concentration was quantified at 48 hours post-infection using the DropArray system (Curiox). All experiments were performed three times (n=3) independently. All bars show mean ± SD with a one-way ANOVA analysis. To correct for multiple testing, the p-value was corrected accordingly (* adjusted p-value<0.0055).

### Neither IL-1β, IL-6 nor LIF protect hiNeurons or hiAstrocytes from Tha-eGFP or Th2P-4M-eGFP infection *in vitro*

Since Tha and Th2P-4M induce the production of IL-1β, IL-6, and LIF in glial cells, we wanted to investigate if these cytokines can restrict RABV replication in hiNeurons or hiAstrocytes via the induction of secondary signalling pathways. Therefore, hiNeurons and hiAstrocytes were infected with Tha-eGFP or Th2P-4M-eGFP and treated with human recombinant IL-1β, IL-6 or LIF two hours after infection. Forty-eight hours post-infection, the percentage of eGFP^+^ hiNeurons and hiAstrocytes was quantified by fluorescence microscopy (Fig 6).

**Fig 6.**
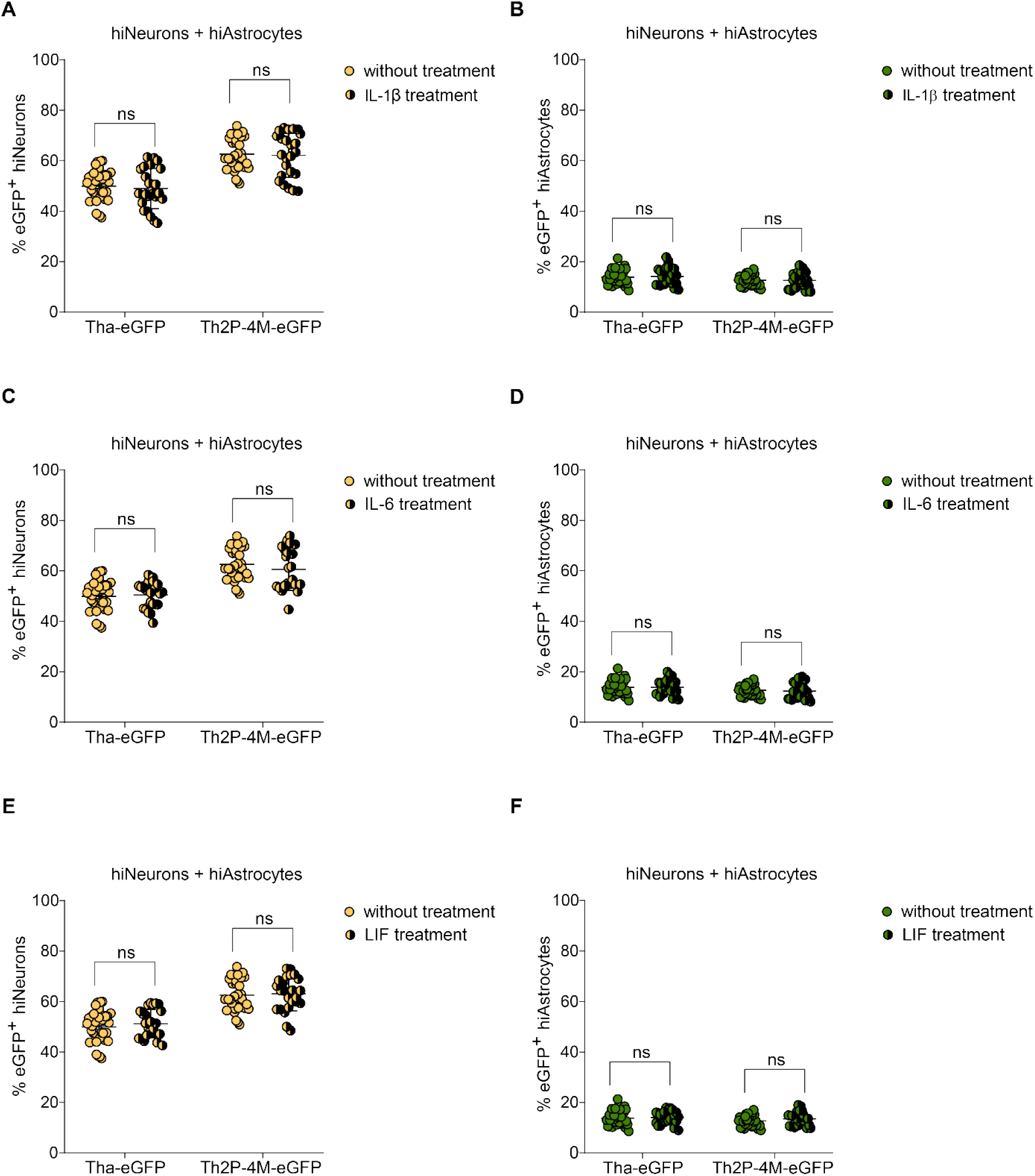
Neither IL-1β, IL-6 nor LIF protect human hiNeurons or hiAstrocytes from Tha-eGFP or Th2P-4M-eGFP infection *in vitro*. (A) Quantification of eGFP^+^ hiNeurons in co-cultures consisting of hiNeurons and hiAstrocytes upon treatment with recombinant human IL-1β. (B) Quantification of eGFP^+^ hiAstrocytes in co-cultures consisting of hiNeurons and hiAstrocytes upon treatment with recombinant human IL-1β. (C) Quantification of eGFP^+^ hiNeurons in co-cultures consisting of hiNeurons and hiAstrocytes upon treatment with recombinant human IL-6. (D) Quantification of eGFP^+^ hiAstrocytes in co-cultures consisting of hiNeurons and hiAstrocytes upon treatment with recombinant human IL-6. (E) Quantification of eGFP^+^ hiNeurons in co-cultures consisting of hiNeurons and hiAstrocytes upon treatment with recombinant human LIF. (F) Quantification of eGFP^+^ hiAstrocytes in co-cultures consisting of hiNeurons and hiAstrocytes upon treatment with recombinant human LIF. (A-F) Prior to infection, cultures consisted of 86% hiNeurons and 14% hiAstrocytes (S3 Table). Cells were infected with Tha-eGFP or Th2P-4M-eGFP (MOI 0.5) and imaged at 48 hours post-infection. Two hours post-infection, the culture medium was removed, and cells were treated with 100 ng/mL recombinant human 1β, IL-6 or LIF. All experiments were performed three times (n=3) independently. Each dot represents imaging of one well of a 96-well-plate (approx. 1×10^4^ cells/well). Bars show mean ± SD with a paired two-tailed t-test (****p-value<0.0001, *** p-value<0.001, ** p-value<0.01, * p-value<0.05). IL-1β = interleukin 1 beta; IL-6 = interleukin 6; LIF = leukemia inhibitory factor.

Fluorescence imaging revealed that neither IL-1β (Fig 6A-B), IL-6 (Fig 6C-D) nor LIF (Fig 6E-F) significantly modulated the percentage of eGFP^+^ hiNeurons or hiAstrocytes at 48 hours post-infection.

## Discussion

Here, we report that cell-dependent factors determine RABV tropism: the simple presence of glial cells in co-cultures as well as the transfer of supernatant from glial cells influences Tha tropism (Fig 2). Additionally, the presence of hiMicros reduces the profound neurotropism of Tha-eGFP (Fig 3B), suggesting a role in microglial-induced communication with neighbouring cells and phagocytosis of debris from infected neurons which was previously shown to be critical in inhibiting viral spread in the CNS (37). Possible underlying mechanisms comprise glial-mediated signalling directly via cell-to-cell contacts or via the secretion of inflammatory proteins (Fig 2 and 3), the induction of inflammatory genes (Fig 4) as well as the induction of cell type-specific inflammatory proteins (Fig 5) which can induce secondary signalling cascades in surrounding cells. Overall, we show that cell type-specific immune responses of distinct CNS cells partly shape RABV tropism *in vitro*.

Apart from cell-dependent factors, virus-intrinsic factors such as conserved residues of viral P- and M-proteins determine RABV tropism: strikingly and in contrast to Tha-eGFP, Th2P-4M-eGFP showed a more profound neurotropism in co-cultures consisting of hiNeurons, hiAstrocytes, and hiMicros (Fig 2C). In contrast, viral titers of immunocompetent HeLa cells were previously shown to be higher upon infection with Tha compared to infection with Th2P-4M at 48 hours post-infection. Nevertheless, the same publication provided evidence that BSR-T7 cells which lack efficient IFN signalling show similar viral titers upon Tha and Th2P-4M infection at 48 hours post-infection (10). Taken together, we suggest that depending on the immunocompetence of the host, accessory functions of viral P- and M-proteins allow viral control of antiviral responses but at the same time negatively influences replication efficiency since viral proteins are needed for viral evasion mechanisms.

Comparing expression of innate immune genes of Tha and Th2P-4M infected cells revealed that Tha specifically evades the neural immune response via P- and M-protein-mediated evasion of NF-κB and JAK-STAT pathways (Fig 4). Although antiviral signalling via IFNs was originally considered as a universal mechanism to control viral infections, recent evidence suggests that neurons lack robust innate immune signalling pathways to minimize their detrimental effects to this non-renewable cell population (38,39). In contrast to neuronal impairment to respond to Tha infection via IFN induction, we show that glial cells induce strong innate immune responses upon Tha and Th2P-4M infection (Fig 4 and 5). Consequently, we suggest that the pronounced neurotropism of Th2P-4M results from its limited capacity to evade strong immune responses, thus restricting cellular tropism to cell types that are more defective in IFN signalling, such as cells of neuronal origin (39). Overall, we provide evidence that RABV neurotropism relies on more than simply the structure of the RABV G-protein (18,40) or the expression of RABV surface receptors (6), but accessory functions of viral P- and M-proteins in viral-mediated immune evasion.

Additionally, we report that Tha-eGFP and Th2P-4M-eGFP infect hiNeurons and hiAstrocytes, whereas hiMicros are not susceptible towards Tha-eGFP nor Th2P-4M-eGFP infection *in vitro* (Fig 3). Although Ray and colleagues previously reported that the tissue culture-adapted ERA strain and mouse-adapted bat SRV strain successfully replicate in primary adult human microglia *in vitro* (25), we suggest that the different nature of RABV strains, their culture adaptation, or insufficient cell purification might account for different susceptibilities towards RABV infection observed between these studies. Consequently, our results question the capability of canine RABV field strains to successfully enter and replicate in human microglia. Microscopic analysis of *post-mortem* human brain tissues of rabid patients revealed a moderate increase in microglia numbers and marked activation of microglia which surrounded degenerated neurons (41) which are referred to as Babes nodules (42) and observed in other viral encephalitides and infectious disorders (43). Given that microglia phagocyte degenerated neurons and thereby take up RABV components (41), viral transcripts that are detected in microglia (44) do not necessarily mean that microglia actively support RABV infection. Nevertheless, more research is needed to elucidate the susceptibility and function of microglia in human rabies infection, particularly by using more sophisticated models which reflect the complexity of the human CNS.

In detail, we identified the upregulation of IFN-inducible genes with antiviral responses (*CXCL10, ISG15, MX1*) in hiMicros as well as an increase in IL-6 protein expression in microglial-like cells upon Tha and Th2P-4M infection. IL-6 is known to exert cellular effects via the anti-inflammatory pathway by binding to the membrane bound IL-6 receptor (IL-6R) or the pro-inflammatory pathway via binding of the soluble form of IL-6R and stimulating a distal response (45). Apart from pathogen defence (46), IL-6 is known to induce pleiotropic effects in tissue regeneration (47,48) and inflammation (49). Further, activated microglia are well-known to release pro-inflammatory cytokines such as IL-6, resulting in cytotoxicity, immune activation, neuronal excitotoxicity and apoptosis (50). Although we show here that IL-6 does not directly restrict RABV replication in hiNeurons or hiAstrocytes (Fig 6), IL-6 might modulate microglia and T-cell activation, BBB permeability (45) and synaptic function (51) during RABV infection. Apart from IL-6, several publications already reported the induction of CXCL10 in microglial cell lines *in vitro* upon RABV inoculation (52,53). In detail, *CXCL10* expression acts during RABV infection as a ‘double-edged sword’: CXCL10 is known to activate glial cells and enhance BBB permeability (54) whereas strong expression causes excessive infiltration of inflammatory cells to the CNS, aggravating RABV pathogenicity (55).

Targeting the highly dynamic activation of microglia state is believed to be a new therapeutic avenue for several neuropathologies (56), including parkinson’s disease (57), depression disorders (58), and acute traumatic brain injury (59). Although neuronal dysfunction rather than disease-associated microglia is believed to cause the fatal outcome of rabies encephalomyelitis (60), targeting the molecular signature of microglia might support RABV clearance from the infected brain.

Like microglia, astrocytes strongly induce the transcription of cytokines and adaptor molecules (*IFIH1, TLR7, IFNB1, CCL5, CXCL10, IL1B, LIF*) upon Tha and Th2P-4M infection *in vitro*. We assume that astrocytes sense virulent Tha via RLRs (*IFIH1, DDX58, DHX58*) and TLRs (*TLR3, TLR7*) in turn inducing the expression of type I IFNs (*IFN-β*), chemokines (*CCL5, CXCL10*), and interleukins (*IL1B*). Both, IFN-β and IL-1β are known to increase BBB permeability, to activate monocytes, microglia, and astrocytes, and to induce the production of neuroprotective mediators (61). Further, astrocytic expression of *CCL5* and *CXCL10* chemokines induces the recruitment of monocytes, macrophages, dendritic cells, T and B lymphocytes, neutrophils, and regulates microglial activity as well as astrocyte survival (61). Taken together, we suggest that astrocytes sense virulent Tha via RLRs and TLRs, triggering the expression of chemokines and interleukins which in turn increase BBB permeability, induce the recruitment and activation of inflammatory and glial cells. Previously, it has been shown that murine astrocytes strongly respond towards infection with a recombinant RABV carrying the G protein of the more neurotropic CVS strain (SAD-G_CVS_) via RLR- and TLRs-induced expression of *IFN-β in vivo*. As a consequence, the IFN response aborts RABV infection of astrocytes (27). In contrast to attenuated RABV strains, a recent quantitative analysis of astrocyte tropism *in vivo* revealed a strong tropism for astrocytes by RABV field strains (8 - 27%). Similarly, we showed that virulent RABV field strain Tha presents a profound astrocyte tropism *in vitro* (CI_95_ 20.71-31.58%, Fig 3C) despite the induction of a strong astrocyte-mediated immune response upon infection (Fig 4 and 5). Nevertheless, considering the mild and non-specific histopathological changes in the CNS of rabies patients (62) as well as viral-mediated inhibition of the BBB opening (63,64), astrocyte-mediated immune responses fail to induce the opening of the BBB and subsequently clear RABV from the infected CNS. Understanding why the astrocyte-induced inflammatory response does not result in the abortion of astrocyte infection in our model might help understanding why astrocytes fail to induce secondary effects such as the opening of the BBB during natural human rabies infection. Thus, more research is necessary to understand the consequences of astrocyte-mediated antiviral responses during RABV infection.

Apart from astrocyte-mediated IFN and interleukin expression, we show that Tha and Th2P-4M induce LIF protein expression in astrocyte-like cells. LIF is a neurotrophic factor known to be released by astrocytes supporting neuronal survival, differentiation, function, and regeneration, as well as oligodendrocyte survival (61). Recently, it was shown LIF expression in astrocytes is among the most differentially regulated genes upon Zika virus infection (65). Considering that LIF is not actively restricting RABV replication in hNSC-derived CNS cultures (Fig 6), we hypothesize that RABV triggers LIF expression in astrocytes to enhance neuronal survival during RABV infection, preserving the intact neuronal network until the final stage of the disease. Although the role of astrocytic LIF expression during RABV infection is not well understood, RABV-mediated modulation of LIF signalling remains a question of significant interest.

## Conclusion

Overall, our results point out that innate immune responses of CNS cell types towards RABV infection *in vitro* strongly vary according to the CNS cell type investigated (Fig 7).

**Fig 7.**
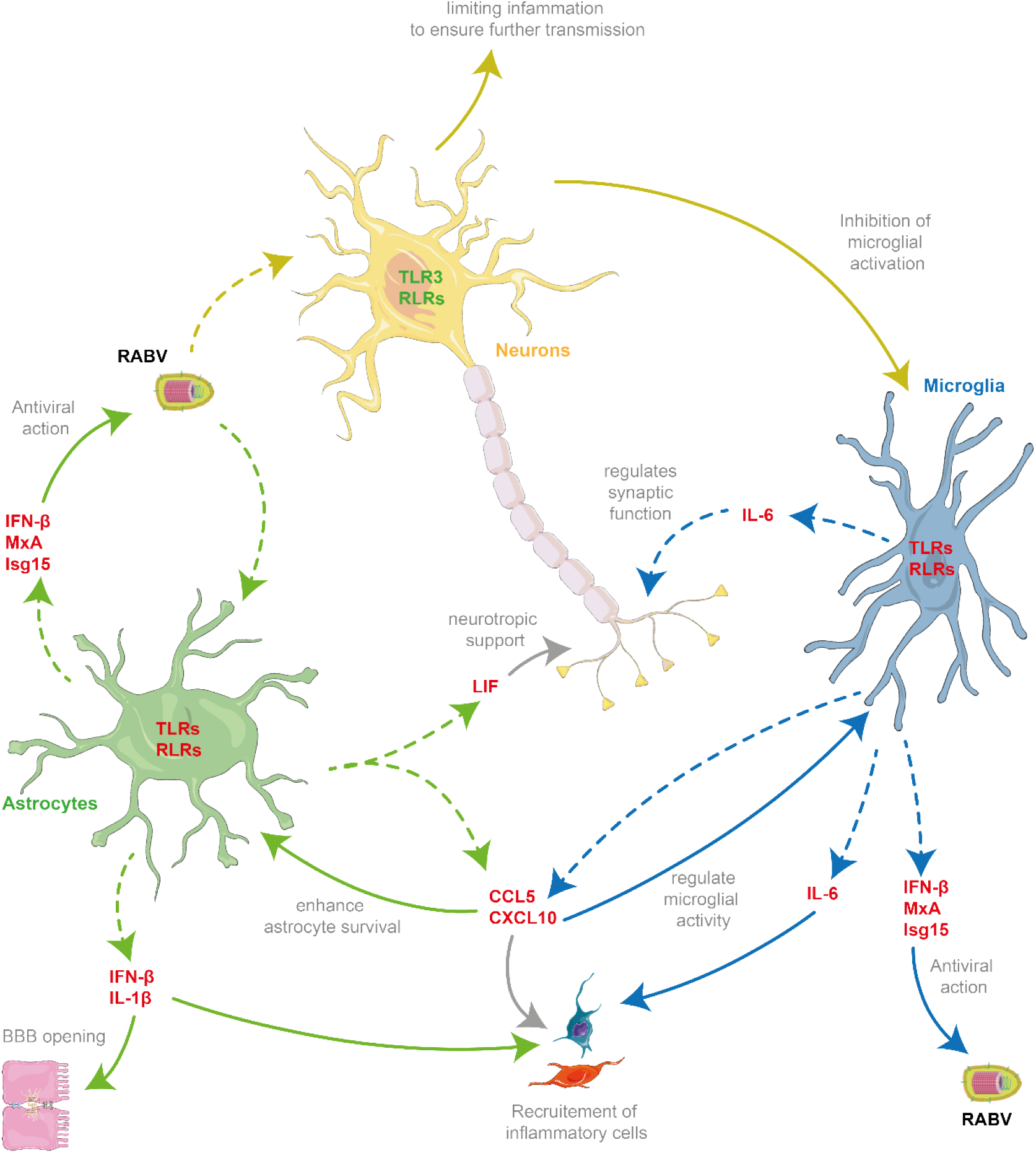
Proposed cellular interactions between neurons, astrocytes and microglia during Tha infection. Whereas hiNeurons and hiAstrocytes were shown to be permissive towards Tha infection, hiMicros are resistant to viral infection *in vitro*. Despite differences in susceptibilities towards virus infection, fetal pAstrocytes and hiMicros mount strong innate immune responses towards Tha infection compared to hiNeurons. Arrows present interactions which were already shown by previous publications. Dashed arrows present genes or proteins which were shown in this study to be upregulated upon virulent Tha infection *in vitro*. Gene or protein names written in green were downregulated during Tha infection, whereas gene names written in red were upregulated during Tha infection.

Whereas neurons lack major antiviral signalling pathways to restrict Tha infection, astrocytes and microglia mount strong immune responses towards Tha infection *in vitro*. Nevertheless, we need to admit that gene and protein expression levels were measured in monocultures which do not reflect the complexity of the CNS. Cellular interactions strongly influence immune responses during physiological and pathological states (66), consequently limiting the validity of our study to extrapolate data directly to the human CNS. Thus, more sophisticated models such as single cell approaches are needed to elucidate the underlying cell type-specific differences in mounting innate immune responses upon RABV infection. Apart from the role of glial cells in restricting RABV replication, we show that RABV tropism strongly depends on the role of accessory functions of viral P- and M-proteins in viral-mediated immune evasion. At last, we conclude that more research is needed to understand the underlying pathways defining RABV tropism and cell type-specific immune responses. Further, the analysis of the distinct responses of infected and non-infected cells as well as the characterization of the transcriptomic profiles of different neural subtypes would help to understand the intimate interplay of the different types and subtypes of cells during RABV infection and how this influences viral tropism.

## Material and methods

### Viruses

Thailand virus, referred to as Tha (isolate 8743THA, EVAg collection Ref-SKU: 014V-02106), is a field strain of RABV isolated from the brain of a Thai patient who died of rabies after being bitten by a rabid dog (EVAg collection, Ref-SKU: 014V-02106) (29). This virus was further adapted to cell culture on BSR cells (a BHK-21 clone, kindly provided by Monique Lafon, Institute Pasteur, Paris) (67). The recombinant Th2P-4M virus harbours the same genetic background as Tha apart from bearing two mutations in the viral P-protein (W265G, M287V) and four mutations in the viral M-protein (R77K, D100A, A104S, M110L) which were previously described (10,31,32). To monitor viral infection, recombinant viral Tha-eGFP and Th2P-4M-eGFP constructs were used which were generated by cloning sequences of eGFP (recovered from pEGFP-C1 plasmid, Promega) into the genetic sequence of Tha (35) and Th2P-4M, respectively. Viral strains were sequenced before used for infection experiments.

### Cell culture and infection

The human neuroblastoma cell line SK-N-SH (ATCC^®^ HTB-11^™^), the human astrocyte-like cell line SVGp12 (ATCC® CRL8621™), and the human microglial cell line HMC3 (ATCC^®^ CRL-3304^™^) were cultured in Dulbecco’s Modified Eagle Medium (10566016, Invitrogen) supplemented with 10% heat-inactivated fetal bovine serum (FCS, S182H-500, Eurobio) at 37°C and 5% CO_2_.

Human neuronal progenitor cells (EnStem-A™ SCC003, Millipore) were cultured on plates previously cultured with geltrex (A1413201, Invitrogen) in complete neural stem cell (CNSC) medium containing Knock-out D-MEM/F-12 (A1370801, Thermo Fisher), 2 mM Glutamax (35050038, Thermo Fisher), 2% StemPro Neural Supplement (A1050801, Thermo Fisher), 20 ng/mL FGF (PHG0026, Thermo Fisher), and 20 ng/mL EGF (PHG0311, Thermo Fisher) for amplification purpose at 37°C and 5% CO_2_.

Human fetal pAstrocytes (N7805100, Thermo Fisher) were cultured on geltrex-coated plates (A1413201, Thermo Fisher) in complete astrocyte medium (CAM) containing D-MEM (10566016, Thermo Fisher), 2% fetal bovine serum (S182H-500, Eurobio) and N-2 Supplement (17502048, Thermo Fisher) at 37°C and 5% CO_2_.

Human iPSCs (XCL-1, IP-001-1V, XCell Science) were cultured on matrigel (354277, Corning)-coated plates in mTeSR1 medium (85850, STEMCELL Technology). The medium was changed daily.

Upon 80% confluency, cells were used for infection experiments. The cell medium was aspirated, and cells were washed once with PBS (10010023, Thermo Scientific) prior to infection. Adjacently, Tha and Th2P-4M were diluted in culture medium according to the appropriate multiplicity of infection (MOI). Cells were incubated with viral suspension for two hours at 37°C and 5% CO_2_. After two hours, the viral suspension was removed, and the appropriate culture medium was added. Twenty-four hours post infection, cells were treated with 2500 U/mL of IFN-α (I4401-100KU, Sigma-Aldrich) and incubated for 24 hours at 37°C and 5% CO_2_ prior to cell lysis.

### Differentiation of hNSC to hiNeurons

To induce neural differentiation, EnStem-A cells were cultured at a density of 5×10^4^ cells/cm^2^ on plates previously coated with geltrex (A1413302, Thermo Fisher) according to the manufacturer’s instructions. One day after seeding, CNSC was changed to neural differentiation medium (NDM) consisting of neurobasal medium (10880022, Thermo Fisher), B-27 Serum Free Supplement (17504044, Thermo Fisher), 2 mM GlutaMAX (35050038, Thermo Fisher), CultureOne Supplement (A33202-01, Thermo Fisher), and 200 *μ*M ascorbid acid (A4403, Sigma-Aldrich). After 21 days of differentiation, differentiated neurons were used for downstream experiments.

### Differentiation of hiPSCs to hiMacs

Human induced macrophage-like cells (hiMacs) were generated from iPSCs according to a published method described by Takata *et al*. (36). In short, 5 ng/mL BMP-4 (314-BP-050, R&D Systems), 50 ng/mL VEGF (293-VE-500, R&D Systems), 2 *μ*M CHIR99021 (4423, TOCRIS) induced mesoderm specification of human iPSC colonies (XCL-1) and hemangioblast-like cell formation (68). Replacement of CHIR99021 by 20 ng/mL FGF-2 (223-FB-500, R&D Systems), followed by maintenance in 15 ng/mL VEGF and 5 ng/mL FGF-2 induced differentiation into the hematopoietic lineage according to a modified protocol from Grigoriadis *et al*. (69). Wnt signalling was inhibited and hematopoietic stem cells were matured by incubation with 50 ng/ml SCF (255-SC-01M), 10 ng/mL FGF-2, 20 ng/mL IL-3 (203-IL-050, R&D Systems), 10 ng/mL IL-6 (206-IL-050, R&D Systems), 10 ng/mL VEGF and 30 ng/mL DKK-1(5439-DK-500) for 6 days and with 50 ng/mL SCF, 10 ng/mL FGF-2, 20 ng/mL IL-3 and 10 ng/mL IL-6 for 4 days. To promote terminal differentiation of hiMacs, cells were cultured with 50 ng/mL CSF-1 (216-MC-01M, R&D Systems) for 14 days (70–72).

### Differentiation of hiMacs to hiMicros

After hiMac generation, hiMacs were positively selected by fluorescence-activated cell sorting (FACS, MoFlo Astrios, Beckman Coulter) using CD45-BV605 (304042, Biolegend, 1:300), CD11b-BV421 (301324, Biolegend, 1:300), CD14-FITC (982502, Biolegend, 1:300), CD163-APC (333610, Biolegend, 1:300) and CX3CR1-PE (341604, Biolegend, 1:300). FACS-sorted cells were co-cultured with 70 – 80% confluent three-weeks old (day 0 = induction of neural induction) hiNeurons for three weeks. After three weeks of co-culture, cells were used for imaging. To obtain a pure culture of hiMicros for downstream quantitative PCR (qPCR) analysis, FACS-sorted hiMacs were co-cultured for three weeks on 24-well-plates in inserts (353495, Falcon, pore size 0.4 *μ*m) with confluent three-weeks old hiNeurons. During co-culture, 50 ng/mL recombinant human CSF-1 (216-MC-010, R & D Systems) and 50 ng/mL human IL-34 (5265-IL-010, R & D Systems) were added every third day to the culture medium.

### Transfer of conditioned medium

Cell lines (SK-N-SH, HMC3, SVGp12) were seeded in 6-well-plates at a density of 1×10^5^ cells/cm^2^. Twenty-four hours after seeding, cells were infected with Tha or Th2P-4M at a MOI of 5. After two hours, the viral suspension was removed, and 1 mL of culture medium was added per well. SK-N-SH cells were seeded into 96-well-plates (655086, Greiner Bio) at a density of 1.8×10^4^ cells/cm^2^. After 24 hours, SK-N-SH previously seeded in 96-well-plates (655086, Greiner Bio) were infected with Tha-eGFP or Th2P-4M-eGFP at a MOI of 0.5. After two hours the viral suspension was removed and replaced via 100 *μ*L of the supernatant taken from previously infected cells (SK-N-SH, HMC3, SVGp12, at 24 hours post-infection) filtered through a 100 kilodalton membrane (28-9322-58, Dutcher). After 24 hours, the medium was removed, and cells were treated with 2500 U/mL of IFN-α (I4401-100KU, Sigma-Aldrich). Cells were imaged at 48 hours post-infection using the Opera Phenix™ High Content Screening System (Perkin Elmer).

### Treatment of hiNeurons with human recombinant IL-1β, IL-6, and LIF

Human NSC were seeded at a density of 15.6×10^4^ cells/cm^2^ into 96-well-plates (655086, Greiner Bio) and differentiated to hiNeurons as described above. Twenty-one days after differentiation, hiNeurons were infected with Tha-eGFP or Th2P-4M-eGFP at a MOI of 0.5. Two hours after infection, the viral suspension was removed and replaced via 100 *μ*L of NDM or NDM supplemented with 100 ng/mL of recombinant human IL-1β (200-01B, Peprotech), IL-6 (200-06, Peprotech), or LIF (300-05, Peprotech). Cells were imaged at 48 hours post-infection using the Opera Phenix™ High Content Screening System (Perkin Elmer).

### Opera Phenix™ High Content Screening Assay

Cell lines were seeded at a density of 1.8×10^4^ cells/cm^2^. In the case of co-cultures, different cell lines were seeded in equal quantities within wells. Twenty-four hours after seeding, cells were infected with Tha-eGFP and Th2P-4M-eGFP. Twenty-four hours post-infection, cells were treated with 2500 U/mL of IFN-α (I4401-100KU, Sigma-Aldrich) and incubated for 24 hours at 37°C and 5% CO_2_ prior to fixation. Cells were fixed using 4% PFA (Alfa Aesar, J61984) for 15 minutes at room temperature, washed with PBS (10010023, Thermo Scientific) and permeabilized using 0.5% triton X-100 (648463, Millipore) for 10 minutes. Cells were stained with primary and secondary antibodies listed in S5 Table according to the manufacturer’s instructions. Apoptotic cells were quantified by the *In situ* cell death detection kit (12156792910, Roche) according to the manufacturer’s instructions. Dead cells were detected with the ReadyProbes® Cell Viability Imaging Kit (R37610, Thermo Fisher). Images were acquired via the Opera Phenix™ High Content Screening System (Perkin Elmer) using the parameters mentioned in S4 Table.

### RNA isolation and cDNA synthesis

Total RNA was isolated by the following procedure: EzDNase (11766051, Thermo Fisher) was used to eliminate genomic DNA. Adjacently, 500 ng or 1 *μ*g of purified RNA was converted into first-strand cDNA using the SuperScript™ VILO™ IV enzyme (11756050, Thermo Scientific) according to the manufacturer’s instructions. For real-time quantitative PCR (RT-qPCR) experiments, cDNA was diluted 1/50 or 1/100, respectively.

### Quantitative PCR

Quantitative PCR, based on the detection of the SYBR Green dye, was performed by using 2.5 *μ*L of the synthesized and diluted cDNA in the presence of 5 *μ*L QuantiTect SYBR Green (204143, Qiagen) and 1 mM specific primers (S6 Table) in a final volume of 10 *μ*L. Oligonucleotides were used for PCR at a concentration of 10 pmol/*μ*L. All the samples were measured in triplicates. Gene expression levels were normalized to the endogenous expression of the housekeeping gene *18S* (Eurofins) and the respective non-infected cells (mock). Quantitative PCR was performed on the 7500 Real-Time PCR System (Applied Biosystem, 7500 Software v2.3) using the following: initial denaturation step (1x repetition, 10 min, 95°C), amplification step (40 repetitions, 15 sec at 95°C, 1 min at 60°C), and melting curve determination step (1x repetition, 15 sec at 95°C, 1 min at 60°C, 15 sec at 95°C, 15 sec at 60°C). The difference in gene expression was calculated as the difference between infected and non-infected samples (ΔΔCT) (73).

### Protein quantification

Cells were seeded at a density of 1×10^5^ cells/cm^2^ and infected with Tha or Th2P-4M at a MOI of 5. Forty-eight hours post-infection, cells were lysed using Procartaplex™ cell lysis buffer (EPX-99999-000, Thermo Scientific) according to the manufacturer’s instructions to quantify protein expression in the cellular cytoplasm. Intracellular protein concentrations of human IFN-β, IFN-γ, IL-1β, IL-6, IL-15, CXCL10, LIF, CCL5, and TNF-α were quantified using a 9-plex Procartaplex assay (PPX-09, Thermo Scientific). In short, DropArray 96-well-plates (96-CC-BD-05, Curiox) were blocked using 1% BSA for 30 min. After blocking, 20 *μ*L of cell lysate was added per well. According to the protocol, plates were stepwise incubated with 5 *μ*L detection antibody, 10 *μ*L streptavidin-PE, and 10 *μ*L reading buffer per well before being read by the Luminex 200™ instrument (Thermo Scientific).

### Statistical analysis

Percentages and means ± standard deviation (SD) were calculated with Prism 9 (Version 9) and R (version 4.0.4). Standard deviation and statistical significance were only calculated when three independent experiments were conducted. Multiple comparisons of data were performed using the GraphPad Prism software (Version 9) with the test being indicated in the figure legend. Principal component analysis and hierarchical clustering on gene expression values was performed using R (version 4.0.4). Heatmaps represent normalized gene expression values that were clustered after centering gene expression values on their mean. Figures were generated using Adobe Illustrator (CC 2019).

## Declarations

### Ethics approval and consent to participate

Not applicable

### Consent for publication

Not applicable

### Availability of data and materials

All data reported in the manuscript are publicly available.

## Authors’ contributions

LF and HB conceived the experimental hypothesis and interpreted the data. LF, FL and HB designed the experiments. LF, GDM and TK performed the experiments. LF and VG analyzed the data. FG provided intellectual guidance. All authors read and approved the final manuscript.

## Acknowledgements

Image acquisition with the Opera Phenix® Plus High Content Screening System (Perkin Elmer) and data analysis via the Columbus software (Perkin Elmer) were conducted with the help of Anne Danckaert and Nathalie Aulner of UTechS Photonic BioImaging (PBI) core at Institute Pasteur Paris. PBI, a member of the national infrastructure France-BioImaging supported by the French National Research Agency (ANR-10-INBS-04), kindly acknowledges the financial support of the Région Ile-de France (program DIM1Health) and the Institute Pasteur.

## Abbreviations

CI_95_: 95% confidence interval
CNS: central nervous system
CNSC: complete neural stem cell
DAMPS: danger-associated molecular patterns
eGFP: enhanced green fluorescent protein
G-protein: glycoprotein
GLAST-1: Glutamate Aspartate Transporter 1
hiMacs: human induced pluripotent stem cells-derived macrophage-like cells
hiMicros: human induced pluripotent stem cell-derived microglia-like cells
hNSC: human neural stem cells
IBA1: Ionized calcium-binding adaptor molecule
IFN-β: interferon beta
IL-1β: interleukin 1 beta
IL-6R: interleukin 6 receptor
iPSC: induced pluripotent stem cells
IRF-3: interferon regulatory factor 3
ISGs: interferon-stimulated genes
JAK: janus kinase
L-protein: RNA-polymerase
LIF: leukemia inhibitory factor
M-protein: matrix protein
mGluR2: metabotropic glutamate receptor subtype 2
MOI: multiplicity of infection
nAChR: nicotinic acetylcholine receptor
NCAM: neuronal cell adhesion molecule
NDM: neural differentiation medium
NF-κB: nuclear factor kappa-light-chain-enhancer of activated B cells
NOD-like receptors: nucleotide-binding oligomerization domain-like receptors
N-protein: nucleoprotein
PAMPS: pathogen-associated molecular patterns
hiAstrocytes: human induced NSC-derived astrocytes
hiNeurons: human induced NSC-derived neurons
P-protein: phosphoprotein
RABV: rabies virus
RIG-I: retinoic acid-inducible gene I
RLRs: retinoic acid-inducible gene I-like helicases
PRRs: pattern recognition receptors
p75NTR: low-affinity p75 neurotropin receptor
RT-qPCR: Real-Time quantitative PCR
STAT: signal transducer and activator of transcription
TLRs: toll-like receptors
TUBB3: Class III Beta-Tubulin

## References

1. Lentz TL, Burrage TG, Smith AL, Crick J, Tignor GH. Is the acetylcholine receptor a rabies virus receptor? Science. 1982;215(4529):182–4.

2. Thoulouze MI, Lafage M, Schachner M, Hartmann U, Cremer H, Lafon M. The neural cell adhesion molecule is a receptor for rabies virus. J Virol. 1998;72(9):7181–90.

3. Tuffereau C, Bénéjean J, Blondel D, Kieffer B, Flamand A. Low-affinity nerve-growth factor receptor (P75NTR) can serve as a receptor for rabies virus. EMBO J. 1998;17(24):7250–9.

4. Wang J, Wang Z, Liu R, Shuai L, Wang X, Luo J, et al. Metabotropic glutamate receptor subtype 2 is a cellular receptor for rabies virus. 2018;2:1–21.

5. Tuffereau C, Schmidt K, Langevin C, Lafay F, Dechant G, Koltzenburg M. The rabies virus glycoprotein receptor p75NTR is not essential for rabies virus infection. J Virol. 2007;81(24):13622–30.

6. Lafon M. Rabies virus receptors. J Neurovirol. 2005;11(1):82–7.

7. Tsiang H, Koulakoff A, Bizzini B, Berwald-Netter Y. Neurotropism of rabies virus. An in vitro study. J Neuropathol Exp Neurol. 1983;42(4):439–52.

8. Potratz M, Zaeck L, Christen M, Kamp V, Klein A, Freuling CM, et al. Astrocyte Infection during Rabies Encephalitis Depends on the Virus Strain and Infection Route as Demonstrated by Novel Quantitative 3D Analysis of Cell Tropism. Cells. 2020;9(1):1– 25.

9. Potratz M, Zaeck LM, Weigel C, Klein A, Freuling CM, Müller T, et al. Neuroglia infection by rabies virus after anterograde virus spread in peripheral neurons. Acta Neuropathol Commun. 2020;8(1):199.

10. Sonthonnax F, Besson BB, Bonnaud E, Jouvion G, Merino D, Larrous F, et al. Lyssavirus matrix protein cooperates with phosphoprotein to modulate the Jak-Stat pathway. Sci Rep. 2019;9(1):1–13.

11. Fooks AR, Banyard AC, Horton DL, Johnson N, Mcelhinney LM, Jackson AC. Current status of rabies and prospects for elimination. Lancet. 2014;384(9951):1389–99.

12. Fooks AR, Cliquet F, Finke S, Freuling C, Hemachudha T, Mani RS, et al. Rabies. Nat Rev Dis Prim. 2017;3:17091.

13. Ugolini G. Rabies virus as a transneuronal tracer of neuronal connections. Adv Virus Res. 2011;79:165–202.

14. Baer GM. The Natural History of Rabies. Vol. 2nd editio, Virus Research. 1991. 640 p.

15. Lafon M. Evasive Strategies in Rabies Virus Infection. 1st ed. Vol. 79, Advances in Virus Research. Elsevier Inc.; 2011. 33–53 p.

16. McFadden G, Mohamed MR, Rahman MM, Bartee E. Cytokine determinants of viral tropism. Vol. 9, Nature Reviews Immunology. 2009. p. 645–55.

17. Akira S, Uematsu S, Takeuchi O. Pathogen recognition and innate immunity. Cell. 2006;124(4):783–801.

18. Etessami R, Conzelmann K-K, Fadai-Ghotbi B, Natelson B, Tsiang H, Ceccaldi P-E. Spread and pathogenic characteristics of a G-deficient rabies virus recombinant: an in vitro and in vivo study. J Gen Virol. 2000;81(Pt 9):2147–53.

19. Le Blanc I, Luyet P-P, Pons V, Ferguson C, Emans N, Petiot A, et al. Endosome-to-cytosol transport of viral nucleocapsids. Nat Cell Biol. 2005;7(7):653–64.

20. Piccinotti S, Kirchhausen T, Whelan SPJ. Uptake of Rabies Virus into Epithelial Cells by Clathrin-Mediated Endocytosis Depends upon Actin. J Virol. 2013;87(21):11637–47.

21. Dietzschold B, Wiktor TJ, Trojanowski JQ, Macfarlan RI, Wunner WH, Torres-Anjel MJ, et al. Differences in cell-to-cell spread of pathogenic and apathogenic rabies virus in vivo and in vitro. J Virol. 1985;56(1):12–8.

22. Wickersham IR, Finke S, Conzelmann KK, Callaway EM. Retrograde neuronal tracing with a deletion-mutant rabies virus. Nat Methods. 2007;4(1):47–9.

23. Wickersham IR, Lyon DC, Barnard RJO, Mori T, Finke S, Conzelmann KK, et al. Monosynaptic Restriction of Transsynaptic Tracing from Single, Genetically Targeted Neurons. Neuron. 2007;53(5):639–47.

24. Prosniak M, Zborek A, Scott GS, Roy A, Phares TW, Koprowski H, et al. Differential expression of growth factors at the cellular level in virus-infected brain. Proc Natl Acad Sci U S A. 2003;100(11):6765–70.

25. Ray NB, Power C, Lynch WPP, Ewalt LCC, Lodmell DLL. Rabies viruses infect primary cultures of murine, feline, and human microglia and astrocytes. Arch Virol. 1997;142(5):1011–9.

26. Sugamata M, Miyazawa M, Mori S, Spangrude GJ, Ewalt LC, Lodmell DL. Paralysis of street rabies virus-infected mice is dependent on T lymphocytes. J Virol. 1992;66(2):1252–60.

27. Pfefferkorn C, Kallfass C, Lienenklaus S, Spanier J, Kalinke U, Rieder M, et al. Abortively Infected Astrocytes Appear To Represent the Main Source of Interferon Beta in the Virus-Infected Brain. J Virol. 2016;90(4):2031–8.

28. AU-Zaeck L, AU-Potratz M, AU-Freuling CM, AU-Müller T, AU-Finke S. High-Resolution 3D Imaging of Rabies Virus Infection in Solvent-Cleared Brain Tissue. JoVE. 2019;(146):e59402.

29. Delmas O, Holmes EC, Talbi C, Larrous F, Dacheux L, Bouchier C, et al. Genomic diversity and evolution of the lyssaviruses. PLoS One. 2008;3(4):1–6.

30. Wiltzer L, Larrous F, Oksayan S, Ito N, Marsh G a., Wang LF, et al. Conservation of a Unique Mechanism of Immune Evasion across the Lyssavirus Genus. J Virol. 2012;86(18):10194–9.

31. Wiltzer L, Okada K, Yamaoka S, Larrous F, Kuusisto HV, Sugiyama M, et al. Interaction of Rabies Virus P-Protein With STAT Proteins is Critical to Lethal Rabies Disease. J Infect Dis. 2014;209(11):1744–53.

32. Ben Khalifa Y, Luco S, Besson B, Sonthonnax F, Archambaud M, Grimes JM, et al. The matrix protein of rabies virus binds to RelAp43 to modulate NF-κB-dependent gene expression related to innate immunity. Sci Rep. 2016;6(9):39420.

33. Besson B, Sonthonnax F, Duchateau M, Ben Khalifa Y, Larrous F, Eun H, et al. Regulation of NF-κB by the p105-ABIN2-TPL2 complex and RelAp43 during rabies virus infection. Schnell MJ, editor. PLoS Pathog. 2017;13(10):e1006697.

34. Luco S, Delmas O, Vidalain P-OO, Tangy F, Weil R, Bourhy H. RelAp43, a Member of the NF-κB Family Involved in Innate Immune Response against Lyssavirus Infection. PLoS Pathog. 2012;8(12):e1003060.

35. Besson B, Kim S, Kim T, Ko Y, Lee S, Larrous F, et al. Kinome-Wide RNA Interference Screening Identifies Mitogen-Activated Protein Kinases and Phosphatidylinositol Metabolism as Key Factors for Rabies Virus Infection. mSphere. 2019;4(3).

36. Takata K, Kozaki T, Lee CZW, Thion MS, Otsuka M, Lim S, et al. Induced-Pluripotent-Stem-Cell-Derived Primitive Macrophages Provide a Platform for Modeling Tissue-Resident Macrophage Differentiation and Function. Immunity. 2017;47(1):183-198.e6.

37. Chhatbar C, Prinz M. The roles of microglia in viral encephalitis: from sensome to therapeutic targeting. Vol. 18, Cellular and Molecular Immunology. Springer Nature; 2021. p. 250–8.

38. Griffin DE. Recovery from viral encephalomyelitis: Immune-mediated noncytolytic virus clearance from neurons. Vol. 47, Immunologic Research. Immunol Res; 2010. p. 123– 33.

39. Yordy B, Iijima N, Huttner A, Leib D, Iwasaki A. A neuron-specific role for autophagy in antiviral defense against herpes simplex virus. Cell Host Microbe. 2012 Sep 13;12(3):334–45.

40. Mebatsion T, Konig M, Conzelmann KK. Budding of rabies virus particles in the absence of the spike glycoprotein. Cell. 1996;84(6):941–51.

41. Liu SQ, Xie Y, Gao X, Wang Q, Zhu WY. Inflammatory response and MAPK and NF-κB pathway activation induced by natural street rabies virus infection in the brain tissues of dogs and humans. Virol J. 2020;17(1):157.

42. Babes V.M. Sur certains caractères des lésions histologiques de la rage. In: Annales de L’Institut Pasteur. 1892. p. 209–23.

43. Love S, Wiley C. Viral diseases. In: Greenfield’s Neuropathology (7th edition). 2002. p. 1–105.

44. Huang KW, Sabatini BL. Single-Cell Analysis of Neuroinflammatory Responses Following Intracranial Injection of G-Deleted Rabies Viruses. Front Cell Neurosci. 2020;14.

45. Rothaug M, Becker-Pauly C, Rose-John S. The role of interleukin-6 signaling in nervous tissue. Vol. 1863, Biochimica et Biophysica Acta - Molecular Cell Research. Elsevier B.V.; 2016. p. 1218–27.

46. Hoge J, Yan I, Jänner N, Schumacher V, Chalaris A, Steinmetz OM, et al. IL-6 Controls the Innate Immune Response against Listeria monocytogenes via Classical IL-6 Signaling . J Immunol. 2013;190(2):703–11.

47. Grivennikov S, Karin E, Terzic J, Mucida D, Yu GY, Vallabhapurapu S, et al. IL-6 and Stat3 Are Required for Survival of Intestinal Epithelial Cells and Development of Colitis-Associated Cancer. Cancer Cell. 2009;15(2):103–13.

48. Matsumoto S, Hara T, Mitsuyama K, Yamamoto M, Tsuruta O, Sata M, et al. Essential Roles of IL-6 Trans -Signaling in Colonic Epithelial Cells, Induced by the IL-6/Soluble– IL-6 Receptor Derived from Lamina Propria Macrophages, on the Development of Colitis-Associated Premalignant Cancer in a Murine Model . J Immunol. 2010;184(3):1543–51.

49. Rabe B, Chalaris A, May U, Waetzig GH, Seegert D, Williams AS, et al. Transgenic blockade of interleukin 6 transsignaling abrogates inflammation. Blood. 2008;111(3):1021–8.

50. Smith JA, Das A, Ray SK, Banik NL. Role of pro-inflammatory cytokines released from microglia in neurodegenerative diseases. Vol. 87, Brain Research Bulletin. Elsevier; 2012. p. 10–20.

51. Gruol DL. IL-6 regulation of synaptic function in the CNS. Neuropharmacology. 2015;96:42–54.

52. Nakamichi K, Saiki M, Sawada M, Takayama-Ito M, Yamamuro Y, Morimoto K, et al. Rabies Virus-Induced Activation of Mitogen-Activated Protein Kinase and NF-B Signaling Pathways Regulates Expression of CXC and CC Chemokine Ligands in Microglia. J Virol. 2005;79(18):11801–12.

53. Zhao P, Yang Y, Feng H, Zhao L, Qin J, Zhang T, et al. Global gene expression changes in BV2 microglial cell line during rabies virus infection. Infect Genet EVol. 2013;20:257–69.

54. Chai Q, She R, Huang Y, Fu ZF. Expression of Neuronal CXCL10 Induced by Rabies Virus Infection Initiates Infiltration of Inflammatory Cells, Production of Chemokines and Cytokines, and Enhancement of Blood-Brain Barrier Permeability. J Virol. 2015;89(1):870–6.

55. Zhao L, Toriumi H, Kuang Y, Chen H, Fu ZF. The roles of chemokines in rabies virus infection: overexpression may not always be beneficial. J Virol. 2009;83(22):11808–18.

56. Fatoba O, Itokazu T, Yamashita T. Microglia as therapeutic target in central nervous system disorders. J Pharmacol Sci. 2020;144(3):102–18.

57. Subramaniam SR, Federoff HJ. Targeting Microglial Activation States as a Therapeutic Avenue in Parkinson’s Disease. Front Aging Neurosci. 2017;9(6):176.

58. Zhang L, Zhang J, You Z. Switching of the Microglial Activation Phenotype Is a Possible Treatment for Depression Disorder. Front Cell Neurosci. 2018;12:306.

59. Chung-Ching C, Mao-Tsun L, Ching-Ping C. Microglial activation as a compelling target for treating acute traumatic brain injury. Curr Med Chem. 2015;22(6):759–70.

60. Fu ZF, Jackson AC. Neuronal dysfunction and death in rabies virus infection. J Neurovirol. 2005;11(1):101–6.

61. Farina C, Aloisi F, Meinl E. Astrocytes are active players in cerebral innate immunity. Vol. 28, Trends in Immunology. Elsevier Current Trends; 2007. p. 138–45.

62. Hemachudha T, Ugolini G, Wacharapluesadee S, Sungkarat W, Shuangshoti S, Laothamatas J. Human rabies: neuropathogenesis, diagnosis, and management. Lancet Neurol. 2013;12(5):498–513.

63. Roy A, Phares TW, Koprowski H, Hooper DC. Failure to open the blood-brain barrier and deliver immune effectors to central nervous system tissues leads to the lethal outcome of silver-haired bat rabies virus infection. J Virol. 2007;81(3):1110–8.

64. Phares TW, Kean RB, Mikheeva T, Hooper DC. Regional Differences in Blood-Brain Barrier Permeability Changes and Inflammation in the Apathogenic Clearance of Virus from the Central Nervous System. J Immunol. 2006;176(12):7666–75.

65. Shereen MA, Bashir N, Su R, Liu F, Wu K, Luo Z, et al. Zika virus dysregulates the expression of astrocytic genes involved in neurodevelopment. Wang R, editor. PLoS Negl Trop Dis. 2021;15(4):e0009362.

66. Armingol E, Officer A, Harismendy O, Lewis NE. Deciphering cell–cell interactions and communication from gene expression. Vol. 22, Nature Reviews Genetics. Nature Research; 2021. p. 71–88.

67. Sato M, Maeda N, Yoshida H, Urade M, Saito S, Miyazaki T, et al. Plaque formation of herpes virus hominis type 2 and rubella virus in variants isolated from the colonies of BHK21/WI-2 cells formed in soft agar. Arch Virol. 1977;53(3):269–73.

68. Kanke K, Masaki H, Saito T, Komiyama Y, Hojo H, Nakauchi H, et al. Stepwise differentiation of pluripotent stem cells into osteoblasts using four small molecules under serum-free and feeder-free conditions. Stem cell reports. 2014;2(6):751–60.

69. Grigoriadis AE, Kennedy M, Bozec A, Brunton F, Stenbeck G, Park I-H, et al. Directed differentiation of hematopoietic precursors and functional osteoclasts from human ES and iPS cells. Blood. 2010;115(14):2769–76.

70. Sturgeon CM, Ditadi A, Awong G, Kennedy M, Keller G. Wnt signaling controls the specification of definitive and primitive hematopoiesis from human pluripotent stem cells. Nat Biotechnol. 2014;32(6):554–61.

71. Ackermann M, Liebhaber S, Klusmann J-H, Lachmann N. Lost in translation: pluripotent stem cell-derived hematopoiesis. EMBO Mol Med. 2015;7(11):1388–402.

72. Chomarat P, Banchereau J, Davoust J, Palucka AK. IL-6 switches the differentiation of monocytes from dendritic cells to macrophages. Nat Immunol. 2000;1(6):510–4.

73. Pfaffl MW. A new mathematical model for relative quantification in real-time RT-PCR. Nucleic Acids Res. 2001;29(9):e45.

